# Temperature-Inducible Precision Guided Sterile Insect Technique

**DOI:** 10.1101/2021.06.14.448312

**Authors:** Nikolay P. Kandul, Junru Liu, Omar S. Akbari

## Abstract

Releases of sterile males are the gold standard for many insect population control programs, and precise sex sorting to remove females prior to male releases is essential to the success of these operations. To advance traditional methods for scaling the generation of sterile males, we previously described a CRISPR-mediated precision-guided sterile insect technique (pgSIT), in which Cas9 and gRNA strains are genetically crossed to generate sterile males for release. While effective at generating F_1_ sterile males, pgSIT requires a genetic cross between the two parental strains which requires maintenance and sexing of two strains in a factory. Therefore, to further advance pgSIT by removing this crossing step, here we describe a next-generation <u>T</u>emperature-<u>I</u>nducible pgSIT (TI-pgSIT) technology and demonstrate its proof-of-concept in *Drosophila melanogaster*. Importantly, we were able to develop a true-breeding strain for TI-pgSIT that eliminates the requirement for sex sorting, a feature that may help further automate production at scale.

## INTRODUCTION

Many insect population control approaches require the generation and release of large numbers of sterile males into natural populations. This control strategy was first proposed in 1955, when Edward Knipling proposed releasing sterile males to suppress insect populations— coined the sterile insect technique (SIT)^1^. SIT has since been successfully implemented to suppress wild populations of a variety of insects^2,3^, such as in the eradication of the new world screw-worm fly, *Cochliomyia hominivorax*, in the U.S. and Mexico^4^. Notwithstanding, Knipling’s vision of sexing sterilized insects to remove females prior to release has been challenging to accomplish, even in the screw-worm example, which has limited the implementation of SIT to other insects.

Finding better ways to sex separate insects is necessary, as field trials and models illustrate that releasing only sterile males significantly improves the efficiency of population suppression and can significantly reduce production costs^1,5^. Furthermore, since females are often the sex that transmit pathogens (e.g. mosquitoes), a reliable sexing method to guarantee female elimination prior to release is highly desirable for the implementation of these programs. Other related methods of insect population control, such as Release of Insects carrying a Dominant Lethal (RIDL)^6^ and the *Wolbachia-mediated* Incompatible Insect Technique (IIT)^7–9^, also require precise sexing methods to avoid female releases. Notably, IIT programs are based on repeated releases of *Wolbachia-infected* males, which are incompatible with wild females that lack the specific *Wolbachia* strain. Even the accidental release of a small fraction of *Wolbachia-infected* fertile females could lead to the wide-scale spread of *Wolbachia*, which would immunize populations against the particular IIT program, underscoring the importance of effective sex separation. However, with a few species-specific exceptions^10,11^, insect sex-sorting can be time consuming, labor intensive, error-prone, and species-dependent ^12–14^.

We recently developed an alternative platform for the generation and sex separation of sterile males using the CRISPR-mediated <u>p</u>recision <u>g</u>uided SIT (pgSIT) technology^15,16^. This technology mechanistically relies on lethal/sterile mosaicism, mediated by the precision and accuracy of CRISPR, to simultaneously disrupt specific genes essential for female viability and male fertility during development, ensuring the exclusive production of sterile males. To generate pgSIT sterile males in this system, two homozygous strains are raised that harbor either Cas9 or guide RNAs (gRNAs), which are genetically crossed to produce F_1_ sterile male progeny that can be deployed at any life stage for population suppression. To further advance this system and mitigate the need for the genetic cross, we herein describe a next-generation <u>T</u>emperature-<u>I</u>nducible pgSIT (TI-pgSIT) technology and demonstrate its proof-of-concept in *Drosophila melanogaster.*

## RESULTS

### Temperature-Inducible Cas9 Activation

To generate an inducible platform that does not require exposure to radiation/chemicals/antibiotics, which can impact the fitness of released animals^17–21^, we utilized a temperature-inducible activation system. We took advantage of the mechanism controlling the expression of *Hsp70Bb*, from the heat-shock 70 family of proteins, which can be temporarily activated by simply raising temperature to 37°C, a heat shock. When the temperature drops, the expression rapidly returns back to pre-shock levels^22–27^. Given this feature, we leveraged the classical Hsp70Bb (Hsp70, Hsp, CG31359) promoter to generate a temperature-inducible Cas9 expression cassette *(Hsp70Bb-Cas9)* (**Supplementary Fig. 1A**). For a visual indicator of promoter activity, we also included a self-cleaving T2A peptide and eGFP coding sequence downstream (3’) from the Hsp-driven Cas9. With this, we established a homozygous transgenic strain of *Drosophila melanogaster.* As the baseline expression of the *Hsp70Bb* promoter at 25°C is well known^28–30^, we compared expression at two temperatures −18°C and 26°C. To visually assess the activity of *Hsp70Bb-Cas9*, we compared GFP fluorescence in *Hsp70Bb-Cas9* embryos, larvae, and adults, raised at either 18°C or 26°C with and without a two-hour heat shock at 37°C (**Fig. 1A**). Without heat shock, we did not detect visible changes in GFP fluorescence between flies raised at 18°C and 26°C (**Fig. 1B**). However, the heat-shocked individuals raised either at 18°C or 26°C had significantly brighter GFP fluorescence, indicating that exposure to 37°C induces robust expression (**Fig. 1B**).

**Figure 1.**
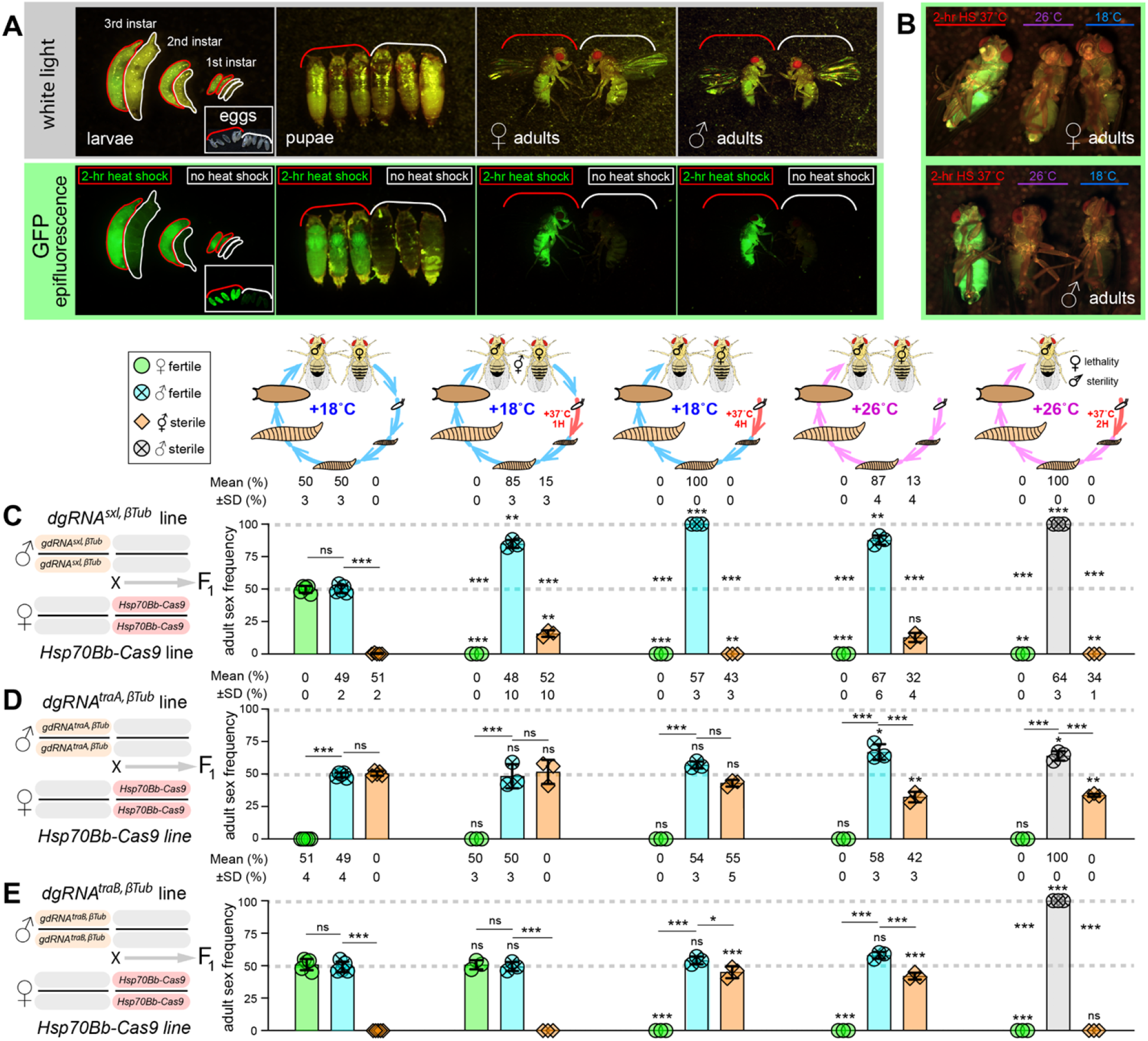
Assessment of temperature inducible pgSIT systems. To establish a visual indicator of Cas9 expression, the GFP coding sequence was attached to the C-terminal end of the *Streptococcus pyogenes*-derived *Cas9* (*Cas9*) coding sequence via a self-cleaving T2A peptide. (**A–B**) A two-hour heat shock at 37°C activates the expression of *Hsp70Bb-Cas9* at the P{CaryP}attP2 site, as indicated by the GFP expression. (**B**) Raising embryos harboring the *Hsp70Bb-Cas9* to adult flies at 26°C does not activate visible GFP fluorescence in living flies. The baseline and activated expression of *Hsp70Bb-Cas9* was tested in combination with three different *dgRNAs:* (**C**) *dgRNA^sxl,βTub^*, (**D**) *dgRNA^traA,βTub^* and (**E**) *dgRNA^traB,βTub^* to assess the feasibility of the <u>T</u>emperature-<u>I</u>nducible <u>p</u>recision <u>g</u>uided <u>S</u>terile <u>I</u>nsect <u>T</u>echnique (TI-pgSIT) design. The staged trans-heterozygous F_1_ embryos generated by reciprocal genetic crosses between homozygous *dgRNAs* and *Hsp70Bb-Cas9* lines were raised at 18°C or 26°C with additional heat shocks at 37°C. The sex and fertility of emerged adult flies was scored and plotted as bar graphs. Since the knockouts of *sxl* and *tra* transform the normal-looking females into intersexes, the emerging F_1_ flies were scored as females (♀), males (♂), or intersexes 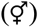. The frequency of each sex that emerged under a specific temperature condition was compared to that of the same sex that emerged at 18°C without a heat shock. Additionally, the male frequency was compared to the female and intersex frequency for each condition. Bar plots show the mean ± standard deviation (SD) over at least three biological replicates. Statistical significance in sex frequency was estimated using a two-sided Student’s *t* test with equal variance. (^ns^*p* ≥ 0.05, \**p* < 0.05, \*\**p* < 0.01, and \*\*\**p* < 0.001).

### Basal Expression of *Cas9*

To genetically determine the basal activity of *Hsp70Bb-Cas9* at 18°C, we performed a series of genetic crosses that would enable us to measure leaky expression. We used constitutively expressing double gRNA (dgRNA) lines that target essential female viability genes, including sex-determination genes *sex lethal* (*sxl*)^31^ or *transformer* (*tra*)^32^ in addition to an essential male fertility gene that is active during spermatogenesis, *βTubulin 85D* (*βTub*)^33^. To target these genes, we used previously generated lines (*dgRNA^sxl,βTub^* and *dgRNA^traA,βTub^*)^15^ and generated a new dgRNA line (*dgRNA^traB,βTub^*) that targets a unique site in *tra*, each constitutively expressing two gRNAs: one targeting *βTub* and one targeting either *sxl* or *tra* (**Supplementary Fig. 1B, Supplementary Table 1**). We crossed homozygous *dgRNA* males to homozygous *Hsp70Bb-Cas9* females and raised the F_1_ progeny at 18°C. The trans-heterozygous F_1_ progeny harboring *Hsp70Bb-Cas9* together with either *dgRNA^sxl,βTub^* or *dgRNA^traB,βTub^* developed into fertile females and males at equal frequencies: 49.8±2.7% ♀ vs 50.1±2.8 ♂ (*p* > 0.884, a two-sided Student’s *t*-test with equal variance; **Fig. 1C, Supplementary. Data 1**), and 51.0±4.1% ♀ vs 49.0±4.1 ♂ (*p* > 0.452, a twosided Student’s *t*-test with equal variance; **Fig. 1E, Supplementary. Data 1**), respectively. Notably, the combination of paternal *dgRNA^traA,βTub^* and maternal *Hsp70Bb-Cas9* resulted in complete conversion of females into intersexes 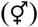 (50.7±1.7% 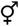 vs 49.3±1.7 ♂, *p* > 0.217, a two-sided Student’s *t*-test with equal variance; **Fig. 1D, Supplementary. Data 1**) suggesting some degree of toxicity likely resulting from the leaky basal activity of *Hsp70Bb-Cas9* combined with *dgRNA^traA,βTub^*. To assess the fertility of the surviving F_1_ progeny from these crosses, we intercrossed F_1_ flies and generated viable F_2_ progeny at 18°C, except from intersex parents which were sterile. The reciprocal genetic cross of *dgRNA* females to *Hsp70Bb-Cas9* males did not cause significant differences in the corresponidng F_1_ sex frequencies (**Supplementary Fig. 2, Supplementary. Data 1**) suggesting that *Hsp70Bb-Cas9* does not induce substantial maternal carryover of Cas9 protein at 18°C. Taken together, these results indicate that the *Hsp70Bb* promoter directed some leaky basal expression sufficient to convert females into intersexes when combined with *dgRNA^traA,βTub^*. However F_1_ trans-heterozygous flies (*dgRNA^sxl,βTub^/+; Hsp70Bb-Cas9/+* and *dgRNA^traB,βTub^/+; Hsp70Bb-Cas9/+*) developed normally into fertile females and males.

Given that generation times in *Drosophila melanogaster* are faster at 26°C, we wanted to also test the possibility of raising trans-heterozygous flies at this temperature. Therefore, we raised trans-heterozygous flies (*dgRNA^sxl,βTub^/+; Hsp70Bb-Cas9/+, dgRNA^traA,βTub^/+; Hsp70Bb-Cas9/+,* and *dgRNA^traB,βTub^/+; Hsp70Bb-Cas9/+*) at 26°C and scored the sex ratios and fertility of emerging flies. Unexpectedly, we found that progeny from these flies could not be maintained at 26°C, since all F_1_ females perished during development, or were converted into sterile intersexes 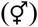, in 12.7±3.5% and 41.9±2.5 of cases, respectively (**Fig. 1C–D**). However, the emerging trans-heterozygous males were fertile, indicating that male sterilization will require additional expression of the CRISPR components (**Fig. 1C–D, Supplementary. Data 1)**. Taken together, these data suggest that the system is sufficiently leaky at 26°C to kill female progeny, yet not leaky enough to sterilize male progeny.

### Temperature-Inducible Phenotypes

To identify the optimal heat-shock conditions required for the complete penetrance of desired TI-pgSIT phenotypes in F_1_ progeny (i.e., female lethality and male sterility), we heat shocked (37°C) F_1_ progeny raised at either 18°C or 26°C and quantified the sex ratios and fertility of emerging progeny. To determine the optimal heat-shock conditions, we compared several conditions. At 18°C, we compared the development with <u>n</u>o <u>h</u>eat <u>s</u>hock (18°C^NHS^); a 1-hr heat shock at the 1^st^ instar larval stage (18°C^1HR-37°C^); or a 4-hr heat shock at the 1^st^ instar larval stage (18°C^4HR-37°C^). At 26°C, we compared the development with <u>n</u>o <u>h</u>eat <u>s</u>hock (26°C^NHS^) or with a 2-hr heat shock at the 1^st^ instar larval stage (26°C^2HR-37°C^) (**Fig. 1C-E, Supplementary Data 1**). The 18°C^1HR-37^° condition killed most of the females expressing *sxl* and transformed the surviving *dgRNA^sxl,βTub^/+; Hsp70Bb-Cas9/+* and *dgRNA^traA,βTub^/+; Hsp70Bb-Cas9/+* transheterozygous females into sterile intersexes (**Fig. 1C–D, Supplementary Data 1**). However this condition was insufficient to transform/kill *dgRNA^traB,βTub^/+; Hsp70Bb-Cas9/+* trans-heterozygous females expressing *U6.3-gRNA^traB^* (**Fig. 1E**). Interestingly, simply increasing the heat-shock period to 4 hours (18°C^4HR-37°C^) completely eliminated the *gRNA^sxl,βTub^/+; Hsp70Bb-Cas9/+* females (**Fig. 1C)** and transformed all *gRNA^traB,βTub^/+*; *Hsp70Bb-Cas9/+* females into intersexes (**Fig. 1E, Supplementary Data 1**). Notwithstanding the complete transformation and killing of females observed above, none of the 18°C^4HR-37°C^, 18°C^1HR-37°C^, and 26°C^NHS^ conditions ensured the complete sterility of F_1_ trans-heterozygous males (**Fig. 1C–E**). Given these results, we next raised trans-heterozygous F_1_ progeny at 26°C with a 2-hr heat shock at the 1^st^ instar larval stage (26°C^2HR-37°C^), which resulted in the development of sterile males and/or sterile intersexes for each trans-heterozygous combination (**Fig. 1C–E, Supplementary Data 1**). Notably, we did not identify *gRNA^traB,βTub^/+*; *Hsp70Bb-Cas9/+* intersex individulas under the 26°C^2HR-37°C^. Taken together, these results indicate that *Hsp70Bb-Cas9* can direct the temperature-inducible expression of Cas9, which is sufficient to cause the 100% penetrance of the desired TI-pgSIT phenotypes. However careful titration is necessary to optimize the temperature conditions to achieve the desired phenotypes.

### Simplified One-Locus TI-pgSIT

Given that both the designed trans-heterozygous combinations generated fertile flies when raised at 18°C and only sterile males when heat shocked (26°C^2HR-37°C^, **Fig. 1C–E**), we next wanted to explore TI-pgSIT systems that function in *cis* to further simplify the approach. Therefore, we engineered two new constructs combining *Hsp70Bb-Cas9* and one of two best dgRNA, *gRNA^sxl,βTub^* and *gRNA^traB,βTub^*, hereafter referred to as *TI-pgSIT^sxl,βTub,Hsp-Cas9^* and *TI-pgSIT^traB,βTub,Hsp-Cas9^,* respectively (**Supplementary Fig. 1C**). Each *TI-pgSIT* cassette was site-specifically inserted into an attP docking site located on the 3rd chromosome (*P{CaryP}attP2*) using φC31-mediated integration^34^ to enable direct comparisons between the two systems. We generated both *TI-pgSIT^sxl,βTub,Hsp-Cas9^* and *TI-pgSIT^traB,βTub,Hsp-Cas9^* transgenic lines and maintained these as heterozygous balanced flies for >10 generations at 18°C. While we were unable to generate a homozygous line for *TI-pgSIT^sxl,βTub,Hsp-Cas9^*, we obtained one for *TI-pgSIT^traB,βTub,Hsp-Cas9^*.

To assess the baseline expression of the one-locus TI-pgSIT systems at 18°C, we evaluated the female-to-male ratio and fertility in lines harboring a copy of either the *TI-pgSIT^sxl,βTub,Hsp-Cas9^* or *TI-pgSIT^traB,βTub,Hsp-Cas9^* cassette. We found a slightly female biased ratio for *TI-pgSIT^sxlβTub,Hsp-Cas9^/+* line maintained at 18°C: 54.5±6.0% ♀ vs 45.5±6.0% ♂ (*p* = 0.025, a two-sided Student’s *t* test with equal variance; **Fig. 2A, Supplementary Data 2**). The *TI-pgSIT^traB,βTub,Hsp-Cas9^* line had a slightly male biased ratio: 47.9±2.8% ♀ vs 52.0±8.3% ♂ for heterozygous flies (*p* < 0.030, a two-sided Student’s *t* test with equal variance; **Fig. 2B, Supplementary Data 2**); and 48.4±2.4% ♀ vs 51.2±2.9% ♂ for homozygous flies (*p* < 0.044, a two-sided Student’s *t* test with equal variance; **Fig. 2C, Supplementary Data 2**). We have maintained both heterozygous *TI-pgSIT^sxl,βTub,Hsp-Cas9^* and homozygous *TI-pgSIT^traB,βTub,Hsp-Cas9^* for nearly 2 years or more than 40 generations and counting at 18°C. Taken together, these experiments indicate that one-locus TI-pgSIT systems can be engineered, expanded, and maintained at 18°C. However, given that we could not generate a homozygous line for *TI-pgSIT^sxl,βTub,Hsp-Cas9^,* again careful titration of expression is necessary.

**Figure 2.**
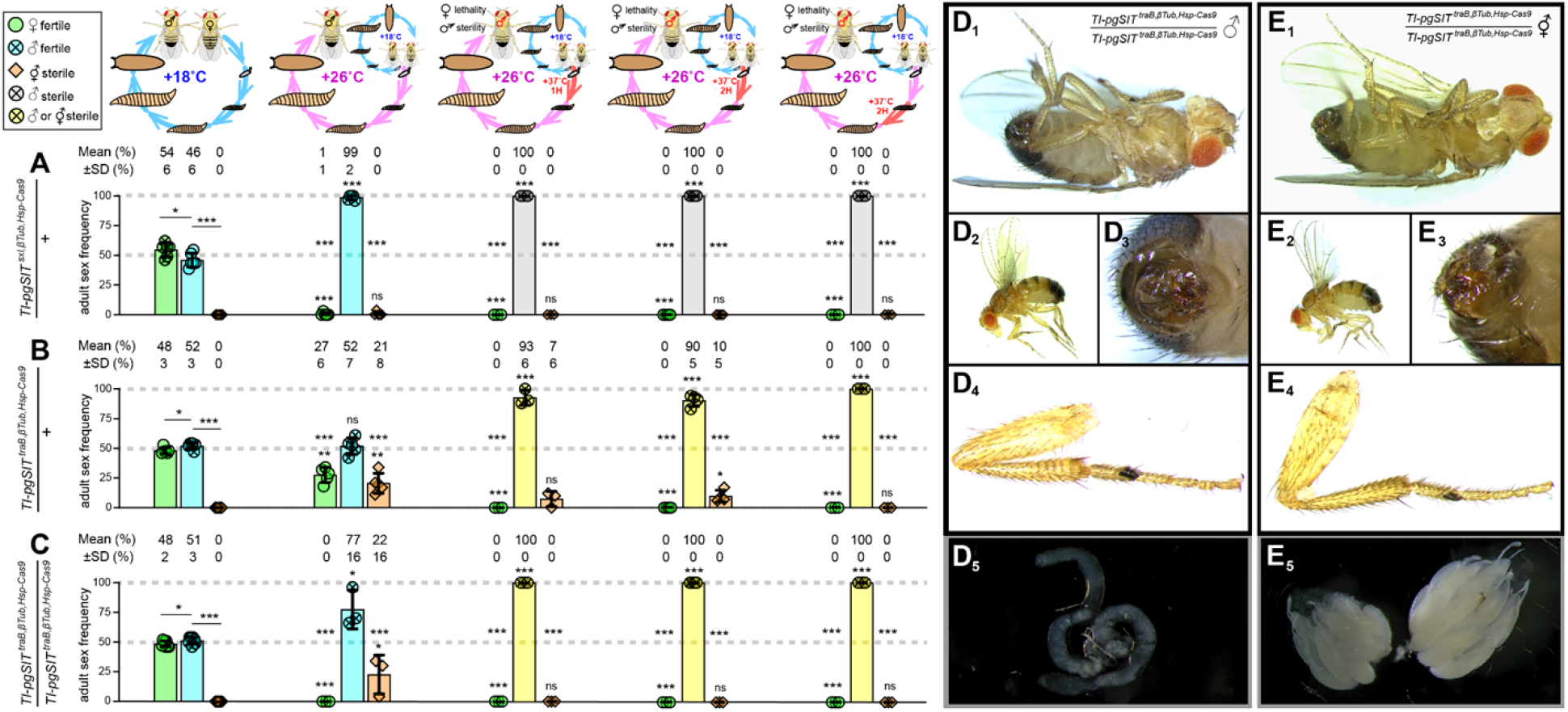
Elevating the temperature of one-locus TI-pgSIT lines produces desired phenotypes. Assessment of phenotypes upon temperature treatments comparing two single locus TI-pgSIT cassettes, (**A**) *TI-dgRNA^sxl,βTub,Hsp-Cas9^* and (**B, C**) *dgRNA^traB,βTub,Hsp-Cas9^*. At 18°C, transgenic flies harboring one or two copies of the TI-pgSIT cassette produce both females and males at a nearly equal sex ratios and can be pure-bred for many generations. The full activation of the TI-pgSIT cassette is achieved by raising the flies at 26°C with an additional heat-shock at 37°C during the first days of development. This activating temperature condition induces 100% penetrance of the pgSIT phenotypes, female-specific lethality and male-specific sterility, and as a result, only sterile males emerge. The sex and fertility of emerged adult flies was scored and plotted as bar graphs. The emerging flies were scored as females (♀), males (♂), or intersexes 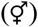. The frequency of each sex that emerged under 18°C treatment was compared to that of the same sex. Additionally, the male frequency was compared to the female and intersex frequency under each condition. Bar plots show the mean ± SD over at least three biological replicates. Statistical significance in sex frequency was estimated using a two-sided Student’s *t* test with equal variance. Pearson’s chi-squared tests for contingency tables were used to assess the difference in male sterility. (^ns^*p* ≥ 0.05, \**p* < 0.05, \*\**p* < 0.01, and \*\*\**p* < 0.001). (**D, E**) Notably, after close examination of heat-shock-induced *dgRNA^traB,βTub,Hsp-Cas9^* males, we inferred that a fraction of flies referred to as males are indeed intersexes. These intersexes have very similar external morphology, including abdomen pigmentation (**E_1-2_**), genitals (**E_3_**), and sex combs (**E_3_**), to that of males (**D_1-4_**) prohibiting their correct identification. Some older intersexes can be identified when, instead of testes (**D_5_**), they develop ovaries (**E_5_**), which result in abdomen extension (**E_2_** vs **D_2_**).

### Temperature-Inducible one-locus TI-pgSIT

We next explored the effects of heat shock on the penetrance of desired TI-pgSIT phenotypes. To activate the *Hsp70Bb-Cas9* expression, we collected eggs from one-locus TI-pgSIT flies maintained at 18°C, and we raised the staged eggs at 26°C with or without an additional heat shock at 37°C. We compared several different heat-shock conditions including: the development from embryos to adult flies at 26°C with <u>n</u>o <u>h</u>eat <u>s</u>hock (26°C^NHS^); with a 1-hr heat shock at the 1st instar larval stage (26°C^1HR-37°C^); or with a 2-hr heat shock at the 1^st^ or 2^nd^ instar larval stages (26°C^2HR-37°C^). For the 26°C^NHS^ condition, when *TI-pgSIT^sxl,βTub,Hsp-Cas9^/+* progeny were raised continuously at 26°C, this resulted in the near-complete elimination of females (54.5±6.0% ♀ at 18°C vs 0.7±1.4% ♀ at 26°C, *p* < 0.0001, a two-sided Student’s *t* test with equal variance) with 98.5±2.0% of males emerging. However not all of these males were sterile (**Fig. 2A, Supplementary Data 2**). Moreover, raising the flies with one or two copies of the *TI-pgSIT^traB,βTub,Hsp-Cas9^* cassette at the 26°C^NSH^ condition affected the sex ratio of the emerging progeny—some or all females, respectively, were transformed into intersexes, though the emerging males were still fertile (**Fig. 2B–C, Supplementary Data 2**). Nevertheless, an additional 1-hr (26°C^1HR-37°C^) or 2-hr (26°C^2HR-37°C^) heat shock of the 1^st^ instar or the 2^nd^ instar larvae harboring either one copy of *TI-pgSIT^sxl,βTub,Hsp-Cas9^* or two copies of *TI-pgSIT^traB,βTub,Hsp-Cas9^* eliminated the females and intersexes and sterilized 100% of the males (**Fig. 2A–C, Supplementary Data 2**). Taken together, these data indicate that heterozygous as well as homozygous viable strains harboring a one-locus *TI-pgSIT* genetic cassette can be generated and maintained at 18°C, and when progeny from these flies are simply grown at 26°C and heat shocked during early larval development, the desired TI-pgSIT is fully penetrant.

### Heat shock induces *TI-pgSIT^traB,βTub,Hsp-Cas9^* intersex flies

We previously observed that *tra* knockout (KO) induces an incomplete masculinization of *D. melanogaster* females converting them into intersexes^15^. To explore further what happens with *TI-pgSIT^traB,βTub,Hsp-Cas9^* females under the 26°C^2HR-37°C^ conditions, heat induced homozygous *TI-pgSIT^traB,βTub,Hsp-Cas9^* males were thoroughly examined. We noticed that several heat shocked TI-pgSIT flies developed extended abdomens. Dissections of their abdomens identified ovaries with oocytes (**Fig. 2E_5_**). Therefore, we inferred that a fraction of *TI-pgSIT^traB,βTub,Hsp-Cas9^* flies, which were raised under the 26°C^2HR-37°C^ conditions, were indeed intersexes. These intersexes, unlike the heat shock induced *dgRNA^traA,βTub^/+; Hsp70Bb-Cas9/+* intersexes reared under the 26°C^2HR-37°C^ (**Fig. 1D**) or the *TI-pgSIT^sxl,βTub,Hsp-Cas9^* intersexes raised under the 26°C without a heat shock (26°C^NHS^, **Fig. 2B–C**), are difficult to distinguish from true males. The abdomen pigmentation (**Fig. 2E_1-2_**), external genitals (**Fig. 2E_3_**), and sex combs (**Fig. 2E_3_**) of the *TI-pgSIT^traB,βTub,Hsp-Cas9^* intersexes reared under 26°C^2HR-37°C^ are nearly identical to those of males (**Fig. 2D_1-4_**) prohibiting their correct identification (**Fig. 2B–C**). Therefore, to avoid intermixing true males with intersexes, we focus on the *TI-pgSIT^sxl,βTub,Hsp-Cas9^* line for further experiments quantifying the basal Cas9 expression and assessing the competitiveness of heat-shocked sterile TI-pgSIT males.

### Fitness and Basal Cas9 Expression

We attempted to establish the homozygous *TI-pgSIT^sxl,βTub,Hsp-Cas9^* line, however homozygous females are only partially fertile and homozygous lineages cannot be maintained. To explore the reasons behind fitness costs of two copies of *TI-pgSIT^sxl,βTub,Hsp-Cas9^* genetic cassette, we examined both *sxl* and *βTub* target sequences in flies raised at 18°C. Using Sanger sequencing, we found that both target sequences were mutagenized resulting in ambiguous sequence reads downstream from the corresponding gRNA cut site (**Supplementary Fig. 3**). These sequencing reads indicate that individual flies were likely mosaic for *wt* and *indel* (i.e. insertion and deletion) alleles at both *sxl* and *βTub* loci. However, it is not clear whether *indel* alleles were induced in only somatic cells or both somatic and germline cells. If functional *indel* alleles, which are resistant to Cas9/dgRNA^*sxl,βTub*^ mediated cleavage, are induced in germline cells they will be selected and propagated through multiple generations. We examined this possibility by assessing the penetrance of heat induced pgSIT phenotypes (aka. female lethality/transformation and male sterility) using both *TI-pgSIT^sxl,βTub,Hsp-Cas9^*/+ and *TI-pgSIT^traB,βTub,Hsp-Casp^/TI-pgSIT^traB,βTub,Hsp-Cas9^* lines after having maintained them for twelve months at 18°C. After heat-shocking multiple batches of larvae and analysing large numbers of flies raised at 26°C, we found that all females either perished or were transformed into intersexes while all resulting males were sterile (**Fig. 3A**). For *TI-pgSIT^sxl,βTub,Hsp-Cas9^/+,* we scored 877 sterile males and a single sterile intersex reared under the 26°C^2HR-37°C^ condition, vs 432 fertile females and 392 fertile males raised at 18°C (**Fig. 3A, Supplementary Data 3**). For *TI-pgSIT^traB,βTub,Hsp-Cas9^/TI-pgSIT^sxl,βTub,Hsp-Cas9^*, we scored 377 sterile males and/or intersexes and no females (**Fig. 2E_1-6_**) following the heat shock (26°C^2HR-37°C^), while 284 fertile females and 274 fertile males emerged under 18°C (**Fig. 3A, Supplementary Data 3**). Taken together, these results suggest that a basal Cas9 expression at 18°C induces some *indel* alleles at *sxl* and *βTub* loci; however the leaky Cas9 expression is likely limited to somatic cells.

**Figure 3.**
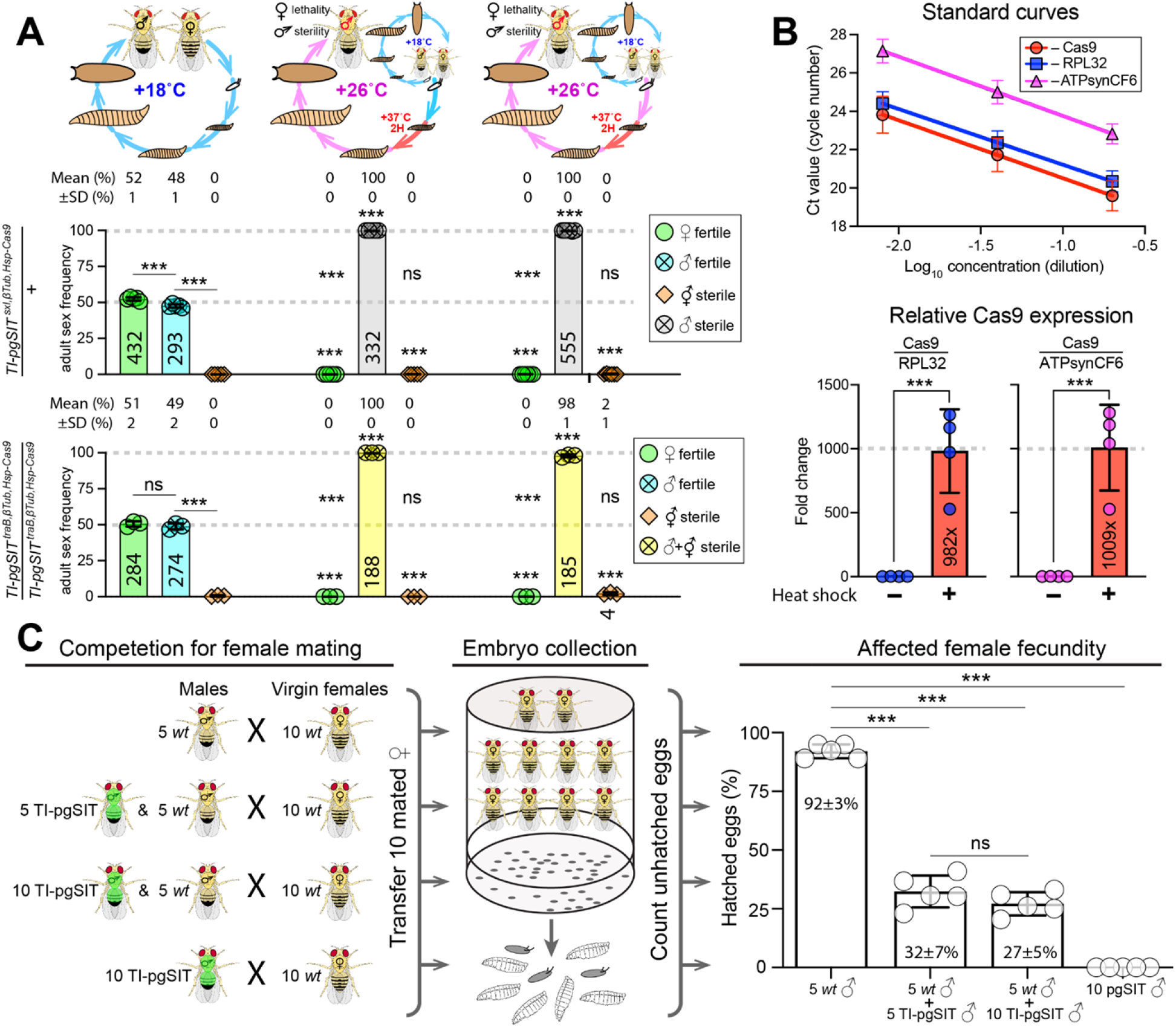
Stability and performance of the TI-pgSIT system twelve months after its development. (**A**) Re-assessment of *TI-pgSIT^sxl,βTub,Hsp-Cas9^* and *TI-pgSIT^traB,βTub,Hsp-Cas9^* one-locus TI-pgSIT lines 12 months later. The sex and fertility of emerged adult flies was scored and plotted as bar graphs. The emerging flies were scored as females (♀), males (♂), or intersexes 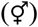, and numbers of scored flies are indicated for each bar. Eggs were collected at 18°C and 26°C, and emerging larvae were heat-shocked at 37°C for 2 hours and then reared at 26°C. The frequency of each sex and its fertility was compared to those of the corresponding sexes reared at 18°C. Additionally, the male frequency was compared to the female and intersex frequency under each condition. Bar plots show the mean ± SD over at least three biological replicates. Statistical significance in sex frequency was estimated using a two-sided Student’s *t* test with equal variance. (^ns^*p* ≥ 0.05, and \*\*\**p* < 0.001). (**B**) In *TI-pgSIT^sxl,β,Tub,Hsp-Cas9^* line, *Cas9* transcription increases nearly 1000 times after a two-hour heat-shock at 37°C. Total RNA was extracted from 2nd instar larvae four hours after heat shock, and RT-qPCR quantification of *Cas9* expression was done relative to *RPL32* and *ATPsynCF6.* (**C**) The heat induced *TI-pgSIT^sxl,βTub,Hsp-Cas9^* males successfully compete against *wt* males to secure matings with *wt* females. The mating success of sterile TI-pgSIT males was evaluated by fecundity decrease (aka. increase of unhatched egg rate). *D. melanogaster* mated female is resistant to the next mating for around 24 hours ^56,57^. Therefore, we confined ten virgin females with multiple males for 12 hours before removing males and assessing female fecundity. The mean percentage of hatched eggs and ± SD over five biological replicates are indicated on the bar graph. We previously showed that a single fertile male was able to fertilize the majority of ten virgin females during 12 hours^15^. The addition of five or ten TI-pgSIT sterile males to five fertile *wt* males resulted in a significant decrease in female fecundity, 92±3% vs 32±7 or 27±5%, respectively. Statistical significance was estimated using a two-sided Student’s *t* test with equal variance. (^ns^*p* ≥ 0.05, and \*\*\**p* < 0.001).

### Quantification of Temperature-Induced Hsp70Bb-Cas9 expression

To assess the extent of temperature induced Cas9 activation, we quantified changes in the Cas9 mRNA relative to other genes using reverse transcription quantitative PCR (RT-qPCR). Two separate constitutively expressed genes, *Ribosomal protein L32* (*RPL32*) and *ATP synthase*, *coupling factor 6* (*ATPsynCF6*), were used for relative quantification of Cas9 expression. We found that a 2-hr 37°C heat shock and 4-hr 26°C incubation of 2^nd^ instar larvae induced a three-order-magnitude increase (1000 times) in the level of the *Cas9* mRNA relative to that in the larvae maintained at 18°C (**Fig. 3B**). Notably, two separate RT-qPCR qualifications based on independent normalizations genes (*RPL32* and *ATPsynCF6*) inferred consistent estimations of the increase in Cas9 expression following the heat shock: 982X as Cas9/*RPL32*, and 1009X as Cas9/*ATPsynCF6* (**Fig. 3B**). Notably, we raised the larvae remaining in the vials and verified that only sterile males emerged from vials raised under the 26°C^2HR-37°C^ condition, while fertile females and males developed in vials maintained at 18°C. Therefore, a single copy of *Hsp70Bb-Cas9* is sufficient to provide a three-order-magnitude transcription increase from its basal expression and induce efficient Cas9/gRNA-mediated mutagenesis, which in turn results in *sxl* and *βTub* knockouts at the organismal level.

### Competitiveness of heat induced *TI-pgSIT^sxl,βTub,Hsp-Cas9^* males

To explore potential fitness costs of activated Cas9 expression, we assessed the competitiveness of heat induced *TI-pgSIT^sxl,βTub,Hsp-Cas9^* males. We previously found that a single pgSIT male generated by crossing *nanos-Cas9* and *dgRNA^sxl,βTub^* were able to court and secure matings with ten *wt* females in the presence of one *wt* males^15^. To further increase competition between males, we confined ten virgin females with five TI-pgSIT or ten TI-pgSIT males in the presence of five *wt* males for 12 hours in the dark (**Fig. 3C**) before removing males and scoring egg hatching rate as female fecundity. Since heat induced TI-pgSIT males are sterile, eggs laid by the *wt* females that mated with TI-pgSIT males will not hatch, and significant decrease in female fecundity will indicate that TI-pgSIT males are able to court, mate, and successfully compete with *wt* males. We found that addition of five or ten TI-pgSIT sterile males to five fertile *wt* males resulted in a significant decrease in female fecundity, 91.9±3.0% vs 32.3±6.8 or 27.1±5.0%, respectively (*P* < 0.0001, a two-sided Student’s *t* test with equal variance, **Fig. 3C**). Interestingly, we did not score a single hatched egg out of 2179 eggs laid by females confined and mated with only TI-pgSIT males (**Fig. 3C, Supplementary Data 4)** further supporting the induced male sterility of TI-pgSIT males. The mating competition assay indicated that the activated *Hsp70Bb-Cas9* expression did not compromise the fitness of TI-pgSIT males and they were highly competitive with *wt* males at courting and mating with *wt* females.

## DISCUSSION

Here we provide the proof-of-concept for a next-generation TI-pgSIT technology. TI-pgSIT addresses two major limitations of the previously described pgSIT^15,16,35^. First, pgSIT relies on the separate inheritance of two required components, Cas9 endonuclease and gRNAs, that are activated in the F_1_ progeny when combined by a genetic cross. As a result, two transgenic lines harboring either the Cas9 endonuclease or gRNAs must be maintained separately, which increases the production costs. Second, though the F_1_ progeny of pgSIT undergo autonomous sex sorting and sterilization during development, enabling their release at any life stage, the genetic cross leading to the production of these F_1_ sterile males requires the precise sex-sorting of parental Cas9 and gRNAs strains. Therefore, although pgSIT ensures the release of only sterile males, it still does not eliminate the insect sex-sorting step. Together these limitations can constrain applications of the original pgSIT technology for insect population control.

The TI-pgSIT system offers possible solutions to these limitations as it instead relies on a single pure-breeding strain, which eliminates the need for maintaining two strains that must still be sex sorted and mated in a facility for production of sterile males. One limitation of the TI-pgSIT approach is the heatshock requirement during F_1_ development, which would preclude the release of eggs. This means that the original pgSIT approach may be better suited for insects with a diapause during the egg stage^15,16^, though both the pgSIT and TI-pgSIT approaches will work well for the insects with a pupal diapause. Other than this limitation, the TI-pgSIT approach retains the benefits of the pgSIT technology, such as its non-invasiveness and high efficiency^15^. Also like the pgSIT approach, TI-pgSIT can in principle be engineered and applied to many insect species with an annotated genome and established transgenesis protocols. It utilizes CRISPR, which works in diverse species from bacteria to humans^36–38^, to disrupt genes that are conserved across insect taxa, such as genes required for sex-determination and fertility. To establish TI-pgSIT in other species, a temperature-inducible promoter is needed. The heat-shock 70 proteins have high interspecies conservation in insects and play important roles in helping them survive under stressful conditions. The *Drosophila Hsp70Bb* promoter is one of the most studied animal promoters^25,26^ and has been widely used for the heat-inducible expression of transgenes in many insect species for over 20 years^39–42^. For example, *Hsp70B* promoters demonstrated robust heat-inducible expression of transgenes in the yellow fever mosquito, *Aedes aegypti*^43^, the Mediterranean fruit fly, *Ceratitis capitata*^44^, and the spotted wing Drosophila, *Drosophila suzukii*^45^. This promoter should therefore be able to drive the heat-inducible expression of Cas9 in many insect species, especially when lower baseline expression is desirable^44^. Moreover, the *Hsp70Bb* promoter could be ideal for inducing positively activated genetic circuits, as the activation of expression is rapid and does not require chemicals or drugs such as antibiotics, which can affect insect fitness directly^17–19^ or indirectly by ablating their microbiomes^20,21^. Unlike common Tet-Off systems with conditional lethal transgenes^6,46,47^ that are derepressed by withholding tetracycline, activation of the *Hsp70Bb* promoter is achieved by elevated temperatures. Heat-shock treatments can reduce maintenance costs compared to other inducible systems, as temperature is relatively costless compared to drugs and antibiotics.

Even though we show that *Cas9* expression can be regulated by temperature using the *Hsp70Bb* promoter, the use of this promoter did result in some leaky expression. The leaky baseline expression of the *Drosophila Hsp70Bb* promoter in somatic cells at 25°C is well known^28–30^ and can be mitigated by either testing multiple genomic integration sites^28^ to titrate the leaky expression, or by targeting different genes. For example, we generated two transgenic lines harboring each TI-pgSIT construct. Flies harboring one or two copies of the *TI-pgSIT^traB,βTub,Hsp-Cas9^* genetic cassette could be pure-bred and maintained at 18°C. However, only one copy of the *TI-pgSIT^sxl,βTub,Hsp-Cas9^* genetic cassette could be maintained at this temperature, as its homozygous female were sterile at 18°C. Because these two lines are inserted at the same genomic insertion site, suggests that the target gene is important. Perhaps the regulation of *sxl* is more sensitive to mosaic mutations in somatic cells than that of *tra*, which would not be surprising as *sxl* is the master gene that controls both female development and X chromosome dosage compensation in *D. melanogaster*, and females homozygous for a loss-of-function mutation died due to the X chromosome hyperactivation ^48^. Nevertheless, a multimerized copy of a Polycomb response element (PRE) could be used to attempt to further suppress the leaky *Hsp70Bb-Cas9* expression^49^ and facilitate homozygousing an engineered TI-pgSIT cassette.

The *Hsp70Bb*-directed expression was reported to be suppressed in germline cells^50^ even in response to heat-shock stimulation^51^. In *Drosophila*, the basic promoter of *Hsp70Bb*, which was incorporated in an upstream activation sequence (UASt) in the Gal4/UAS two-component activation system^52^, was shown to be targeted by Piwi-interacting RNAs (piRNAs) in female germline cells leading to degradation of any mRNA harboring endogenous *Hsp70Bb* gene sequences^53^. We inferred the presence of *indel* alleles at both *sxl* and *βTub* target sites by Sanger sequencing these loci in *TI-pgSIT^sxl,βTub,Hsp-Cas9^* flies raised at 18°C. After maintaining *TI-pgSIT^sxl,βTub,Hsp-Cas9^/+* and *TI-pgSIT^traB,βTub,Hsp-Cas9^TI-pgSIT^traB,βTub,Hsp-Cas9^* lines for twelve months at 18°C, we re-confirmed the complete penetrance of heat induced pgSIT phenotypes (aka. female lethality/transformation and male sterility). Taken together, our results are consistent with the absence of basal Cas9 expression in germline cells. The piRNA-mediated degradation of mRNA molecules harboring *Hsp70Bb* sequences in germline cells safeguards against generation and accumulation of mutant alleles. However, the three-order-magnitude activation (1000x) of Cas9 expression following the heat shock ensures the complete penetrance of both pgSIT phenotypes at the organismic level without compromising male mating competitiveness.

In summary, here we demonstrate that by using a temperature-inducible CRISPR based approach, we can maintain a single true-breeding strain and induce the production of sterile and competitive males simply by shifting the temperature. This opens an entirely new approach for the generation of sterile males, and has now completely eliminated the need for sex sorting that is still required by other similar methods. In the future, TI-pgSIT could be adapted to both agricultural pests and human disease vectors to help increase the production of food and reduce human disease, respectively, thereby eliminating the need for harmful insecticides and revolutionizing insect population control.

## METHODS

### Assembly of genetic constructs

All genetic constructs generated in this study were engineered using Gibson enzymatic assembly^54^. To assemble *Hsp70Bb-Cas9^dsRed^* (**Supplementary Fig. 1A**), the *BicC-Cas9* plasmid^55^ was digested with NotI and PmeI to remove the *BicC* promoter. The 476-base-long fragment encompassing the *Hsp70Bb* promoter and cloning overhangs were PCR amplified from the pCaSpeR-hs plasmid (GenBank #U59056.1) using primers 1137.C1F and 1137.C3R and cloned inside the linearized plasmid (**Supplementary Table 1**). Then, the *Hsp70Bb-Cas9-T2A-eGFP-p10* fragment was subcloned from *Hsp70Bb-Cas9^dsRed^* into the mini-*white* plasmid with the attB site. The *dgRNA^TraB,βTub^* plasmid was assembled following the strategy used to build *dgRNA^Sxl,βTub^* in a previous work^15^ (**Supplementary Fig. 1B**). Briefly, the *U6.3-gRNA^TraB^* fragment was PCR amplified from the *sgRNA^Tra-B^* plasmid using primers 2XgRNA-5F and 2XgRNA-6R and was cloned into the *sgRNA^βTub^* plasmid (Addgene #112691). To build the *TI-pgSIT^sxl,βTub,Hsp-Cas9^* and *TI-pgSIT^TraB,βTub,Hsp-Cas9^* constructs (**Supplementary Fig. 1C**), the U6.3 3’-UTR fragment was amplified using primers 1098A.C1F and 1098A.C2R from the pVG185_w2-y1 plasmid (GenBank #MN551090.1)^55^ and the *Hsp70Bb-Cas9-T2A-eGFP-10* fragment was amplified using primers 1098A.C3F and 1098A.C6R from the *Hsp70Bb-Cas9* plasmid. Both were cloned into the *dgRNA^Sxl,βTub^* (Addgene #112692) or *dgRNA^TraB,βTub^* plasmid, respectively, after linearization at XbaI. The gRNA and primer sequences used to assemble the genetic constructs in the study are presented in **Supplementary Table 1**.

### Fly transgenesis

Embryo injections were carried out at Rainbow Transgenic Flies, Inc. (http://www.rainbowgene.com). We used φC31-mediated integration^34^ to insert the *Hsp70Bb-Cas9^dsRed^* construct at the PBac{y+-attP-3B}KV00033 site on the 3^rd^ chromosome (BDSC #9750) and to insert the *Hsp70Bb-Cas9^dsRed^* construct at the P{CaryP}attP2 site on the 3rd chromosome (BDSC # 8622). The *dgRNA^TraB,βTub^* construct was inserted at the P{CaryP}attP1 site on the 2nd chromosome (BDSC # 8621), and the *TI-pgSIT^traB,βTub,Hsp-Cas9^* and *TI-pgSIT^traB,βTub,Hsp-Cas9^* constructs were inserted at the P{CaryP}attP2 site on the 3rd chromosome (BDSC # 8622). We maintained the embryos injected with the *TI-pgSIT^sxl,βTub,Hsp-Cas9^* and *TI-pgSIT^traB,β,Hsp-Cas9^* constructs and any of their progeny starting from the G_1_ generation at 18°C. Recovered transgenic lines were balanced on the 2^nd^ and 3^rd^ chromosomes using single-chromosome balancer lines (*w^1118^*; *CyO/sna^Sco^* for II and *w^1118^*; *TM3, Sb^1^/TM6B, Tb^1^* for III).

### Fly maintenance and genetics

Flies were examined, scored, and imaged on the Leica M165FC fluorescent stereo microscope equipped with the Leica DMC2900 camera. We tracked the inheritance of *Hsp70Bb-Cas9^dsRed^* using the *Opie2-dsRed* genetic marker. The other transgenes were tracked using the mini-*white* marker. All genetic crosses were performed in the *w-* genetic background. Flies harboring both *Hsp70Bb-Cas9* and *dgRNAs* in the same genetic background were maintained at 18°C with a 12H/12H light and dark cycle, while the flies harboring either *Hsp70Bb-Cas9* or *dgRNAs* were raised under standard conditions at 26°C. All genetic crosses were performed in fly vials using groups of seven to ten flies of each sex.

We first assessed the heat–shock-induced activation of *Hsp70Bb-Cas9* by visualizing GFP fluorescence. The GFP coding sequence was attached to the C-terminal end of the *Streptococcus pyogenes*-derived *Cas9* (*SpCas9*) coding sequence via a self-cleaving T2A peptide and served as a visual indicator of Cas9 expression. The embryos that were laid overnight as well as the larvae, pupae, and adult flies of both *Hsp70Bb-Cas9* and *TI-pgSIT^traB,βTub,Hsp-Cas9^* females or males rearedhomozygous lines were heat shocked for two hours at 37°C, and in 6, 15, or 24 hours post heat shock, their GFP expression was imaged and compared to that of the nontreated embryos, larvae, pupae, or flies raised at 18°C or 26°C. To assess the inducible expression of *Hsp70Bb-Cas9* directly, we compared the Cas9/dgRNA knockout phenotypes induced by a heat shock to those without the heat shock. We tested two different double guide RNA (dgRNA) (*dgRNA^sxl,βTub^* and *dgRNA^traB,βTub^*) lines with the same *Hsp70Bb-Cas9* line as the F_1_ trans-heterozygotes—the classic pgSIT. The homozygous dgRNA and Cas9 lines were genetically crossed, and their trans-heterozygous embryos were raised at either 18°C or 26°C. Additionally, groups of these embryos underwent various durations of heat shocks at 37°C during the 1^st^ or 2^nd^ day post oviposition (**Fig. 1**). For heat-shock treatments, glass vials with staged embryos and/or larvae were incubated in a water bath at 37°C. We tested different temperature conditions to assess the induction levels between the baseline and complete expression of Cas9 for each dgRNA construct: the development at 18°C with <u>n</u>o <u>h</u>eat <u>s</u>hock (18°C^NHS^), a 1-hr heat shock at the 1^st^ instar larval stage (18°C^1HR-37°C^), or a 4-hr heat shock at the 1^st^ instar larval stage (18°C^4HR-37°C^).The development at 26°C was tested with <u>n</u>o <u>h</u>eat <u>s</u>hock (26°C^NHS^) or with a 2-hr heat shock at the 1^st^ instar larval stage (26°c^2HR-37°C^) (**Fig. 1**).

The generated transgenic lines harboring one or two copies of *TI-pgSIT^sxl,βTub,Hsp-Cas9^* and *TI-pgSIT^traB,βTub,Hsp-Cas9^* genetic cassettes were maintained for >10 generations at 18°C. To induce the pgSIT phenotypes, staged embryos were generated at 18°C and shifted to 26°C to complete their development. We assessed different temperature conditions to fully activate the Cas9 expression: the development at 18°C with <u>n</u>o <u>h</u>eat <u>s</u>hock (18°C^NHS^) and the development at 26°C with <u>n</u>o <u>h</u>eat <u>s</u>hock (26°C^NHS^), a 1-hr heat shock at the 1^st^ instar larval stage (26°C^1HR-37°C^), or a 2-hr heat shock at the 1^st^ or 2^nd^ larval stages (26°C^2HR-37°C^) (**Fig. 2**). To estimate the efficiency of knockout phenotypes, we scored the sex of emerging adult flies as female (♀), male (♂), or intersex 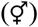 and tested the fertility of generated flies as previously described in Kandul et al.^15^. Note that the induced male sterility was tested in multiple groups of 7–20 males per group from the same biological sample. A single fertile male would designate an entire sample as fertile. Each experimental test was repeated a minimum of three times for statistical comparisons.

### Genotyping loci targeted with gRNAs

We examined the molecular changes that caused female lethality and male sterility following the previously described protocol^15^. Briefly, the *sxl*, *tra*, and *βTub* loci targeted by the gRNAs were PCR amplified from individual flies and were sequenced in both directions using the Sanger method at GeneWiz®. The sequence reads were aligned against the corresponding reference sequences in SnapGene® 4. The primer sequences used for the PCR of the *sxl*, *tra*, *βTub* loci are presented in **Supplementary Table 1**. We also sequenced *sxl* and *βTub* loci using DNA extracted from multiple *TI-pgSIT^sxl,βTub,Hsp-Cas9^* females or males reared at 18°C to assess leaky *Hsp70Bb-Cas9* expression in somatic cells.

### Reverse transcription quantitative PCR (RT-qPCR)

We used the *TI-pgSIT^sxl,βTub,Hsp-Cas9^* line to quantify the activation *Hsp70Bb-Cas9* expression. Vials containing staged larvae were maintained at 18°C. Heat treated vials were incubated in the heat block for 2 hours at 37°C and then for 4 hours at 26°C. The not-heat-treated vials stayed at 18°C. Larvae were separated from food in room-temperature water. Total RNA was extracted using the RNeasy Mini Kit (Qiagen), quantified using the NanoDrop 2000 (Thermo Scientific™), and then treated with DNAse I (Thermo Scientific™) following the protocol. cDNA was synthesized using the RevertAid First Strand cDNA Synthesis Kit (Thermo Scientific™) with a primer mixture of 1:6 of Oligo (dT)_18_ primer and random hexamer primers. Real-time qPCR was performed using LightCycler® 96 Instrument (Roche). RT-qPCR quantification of *Hsp70Bb-Cas9* expression was done relative to *RPL32* and *ATPsynCF6*. Reversed transcribed cDNA samples from not-heat-treated replicates were serially diluted over 50x to build standard curves for each amplified gene fragment and test primer performance (**Supplementary Table 1**). A 10x dilution of cDNA (middle of the standard curve range) was used for relative quantification of *Hsp70Bb-Cas9* expression. Real-time quantification PCR reactions (20μL) contained 4μl of sample, 10μl of SYBR Green Master Mix, 0.8μl of forward primer and 0.8μl of reverse primer and 4.4μl of ultrapure water. Negative control (20μl) contains 10μl of SYBR Green Master Mix, 0.8μl of forward primer and 0.8μl of reverse primer and 8.4 μl of ultrapure water. Three technical replicates were run per place for each of four biological replicates. The Real-time quantification PCR data were analyzed in LightCycler® 96 Application (Roche Applied Science) and exported into an Excel datasheet for further analysis. RNA levels were normalized to *RPL32* or *ATPsynCF6* to generate two separate relative quantifications of *Hsp70Bb-Cas9* mRNA after a two-hour heat shock.

### Competition assay of TI-pgSIT males

We evaluate the competitiveness of the induced *TI-pgSIT^sxl,βTub,Hsp-Cas9^* males by their ability to mate with females in the presence of *wt* males. We previously demonstrated that one fertile male is able to mate with the majority of ten virgin females in twelve hours^15^. To increase mating competition, we confined 10 virgin females with 5 *wt* males alone, 5 *wt* and 5 TI-pgSIT males, 5 *wt* and 10 TI-pgSIT males, or 10 TI-pgSIT males alone in a vial for 12 hours in the dark. As previously, freshly emerged induced TI-pgSIT and *wt* males were isolated from females and aged for 4 days before the competition assay to increase the male courtship drive. After 12 hours of mating, the females were transferred into small embryo collection cages (Genesee Scientific 59–100) with grape juice agar plates and percentage of hatched eggs were calculated. The decrease in female fecundity, estimated by a number of unhatched eggs, indicated the ability of a sterile TI-pgSIT male to score successful mating with females in the presence of a *wt* male; and thus provided a readout of the competitiveness of the induced *TI-pgSIT^sxl,βTub,Hsp-Cas9^* males.

### Statistical analysis

Statistical analyses were performed in JMP 8.0.2 by SAS Institute Inc. Three to five biological replicates were used to generate statistical means for comparisons. *P* values were calculated for a two-sided Student’s *t* test with equal variance.

## DATA AVAILABILITY

All data underlying **Figs. 1–2** are represented fully within **Supplementary Data 1–3**. The plasmids constructed in the study were deposited at Addgene.org (#149424–149427, **Supplementary Fig. 1**). The *Hsp70Bb-Cas9, Hsp70Bb-Cas9^dsRed^, dgRNA^TraB,βTub^, TI-pgSIT^sxl,βTub,Hsp-Cas9^,* and *TI-pgSIT^traB,βTub,Hsp-Cas9^* transgenic lines were deposited to the Bloomington Drosophila Stock Center.

## AUTHOR CONTRIBUTIONS

O.S.A and N.P.K. conceived the concept. J.L. engineered plasmids and performed all molecular work. N.P.K. performed all genetic experiments. All authors analyzed the data, contributed to the writing of the manuscript, and approved the final manuscript.

## ACKNOWLEDGMENTS

This work was supported in part by an NIH RO1 (1R01AI151004-01) award and a (DARPA) Safe Genes Program Grant (HR0011-17-2-0047) both awarded to O.S.A.

## ASSOCIATED CONTENT

Supplementary information is available for this paper at https://doi.org/xxxxx.

---

**Supplementary Figure 1.**
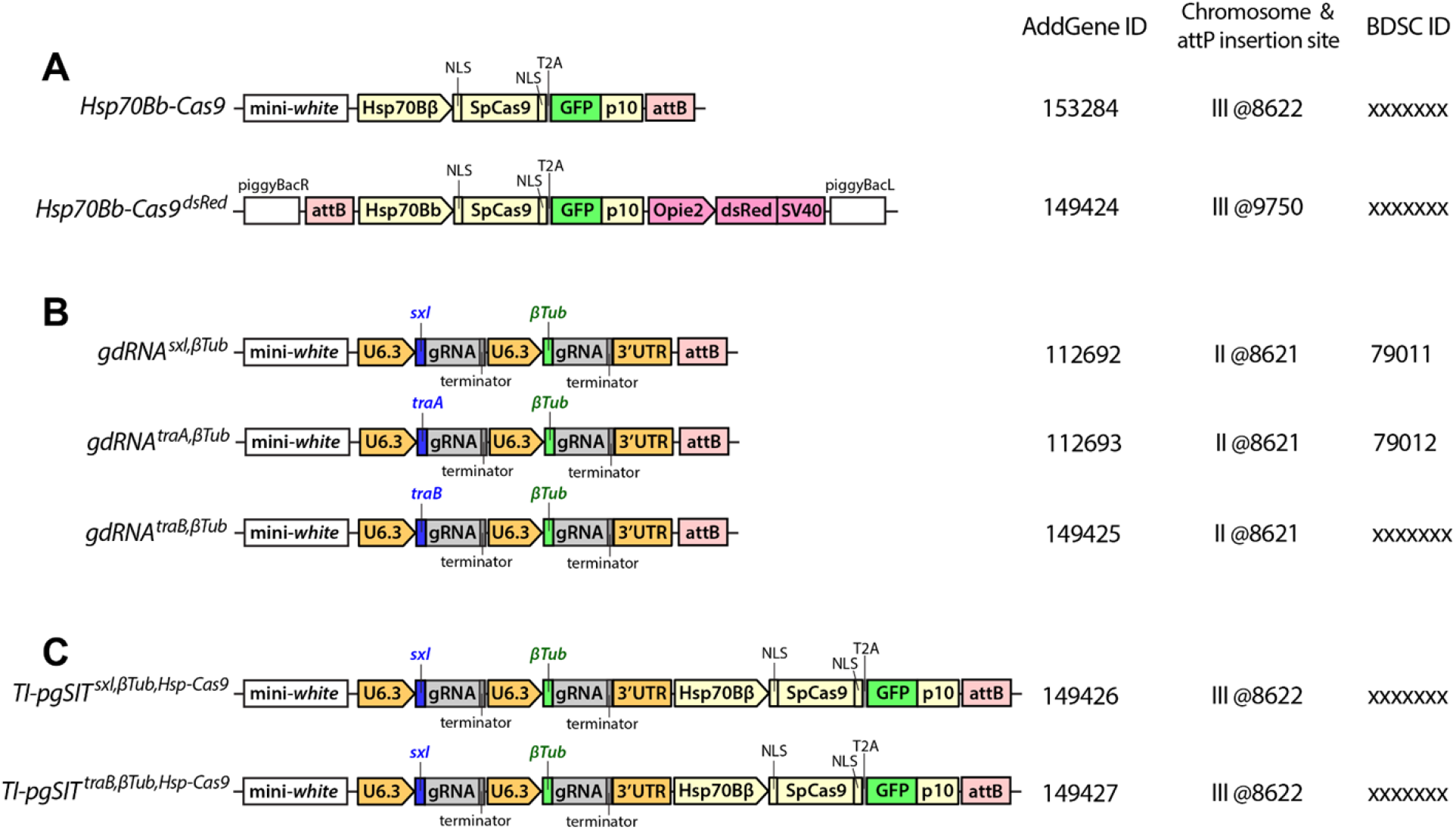
Schematic of genetic constructs used in this study. (**A**) The *Drosophila heatshock protein 70B* (*Hsp70Bb*) promoter directs the temperature-inducible expression of Cas9. The coding sequence of the *Streptococcus pyogenes-derived Cas9* (*Cas9*) was flanked by two nuclear localization signals (NLS) at both ends, to promote nuclear localization, and a self-cleaving T2A peptide with GFP coding sequence at the C-terminal end, serving as a visual indicator of Cas9 expression. The *Opie2-dsRed-SV40* marker transgene was included in the *Hsp70Bb-Casal9* constructs. (**B**) Double guide RNA (dgRNA) genetic constructs. The constitutive expression of two gRNAs targeting *βTubulin 85D* (*βTub*) and either *sex lethal* (*sxl*) or *transformer* (*tra*) is achieved by the *Drosophila* U6.3 promoter. The dgRNA constructs are tracked by the mini-*white* marker gene. The gRNA sequences are indicated in the **Supplementary Table 1**. (**C**) <u>T</u>emperature-<u>I</u>nducible <u>p</u>recision guided <u>S</u>terile <u>I</u>nsect <u>T</u>echnique (TI-pgSIT) genetic cassettes. The *Hsp-70Bb-Cas9-T2A-GFP-p10* fragment was added to the two dgRNA constructs to build two TI-pgSIT cassettes. The genetic cassettes were site-specifically integrated at the *P{CaryP}attP2* site on the 3rd chromosome (BDSC #8622). The genetic constructs and *Drosophila* transgenic lines generated in the study were deposited at Addgene.org and the Bloomington Drosophila Stock Center.

**Supplementary Figure 2.**
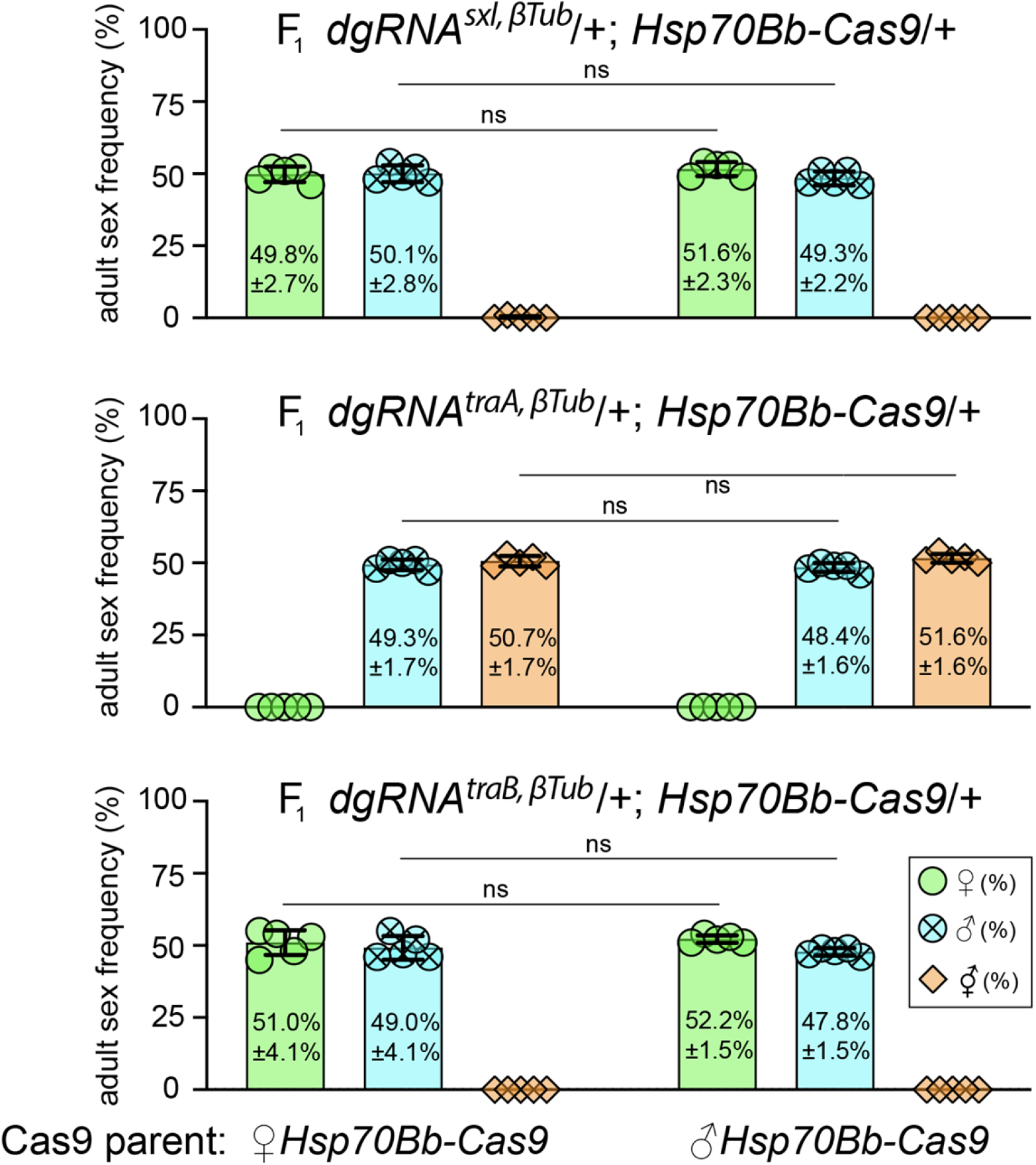
At 18°C, Cas9 protein carryover induced by maternal *Hsp70Bb-Cas9* does not affect F_1_ sex frequenies. To explore whether a leaky Cas9 expression under 18°C causes the maternal protein carryover affecting the progeny sex frequencies, we genetically crossed homozygous *Hsp70Bb-Cas9* line to each of three homozygous dgRNA lines in both directions and compared sex frequencies of F_1_ trans-heterozygotes harboring Cas9 inherited from mothers (maternal Cas9, *Hsp70Bb-Cas9* ♀ x *dgRNA* ♂) or fathers (paternal Cas9, *Hsp70Bb-Cas9* ♂ x *dgRNA* ♀). We did not identified significant differences in sex frequencies between progenies harboring maternal vs paternal Cas9 reared under 18°C (**Supplementary Data 1**). These findings suggest that the basal expression of *Hsp70Bb-Cas9* does not cause the Cas9 maternal carryover affecting the progeny sex frequencies. Bar plots show the mean ± SD over five biological replicates. Statistical significance in sex frequency was estimated using a two-sided Student’s *t* test with equal variance. (^ns^*p* ≥ 0.05).

**Supplementary Figure 3.**
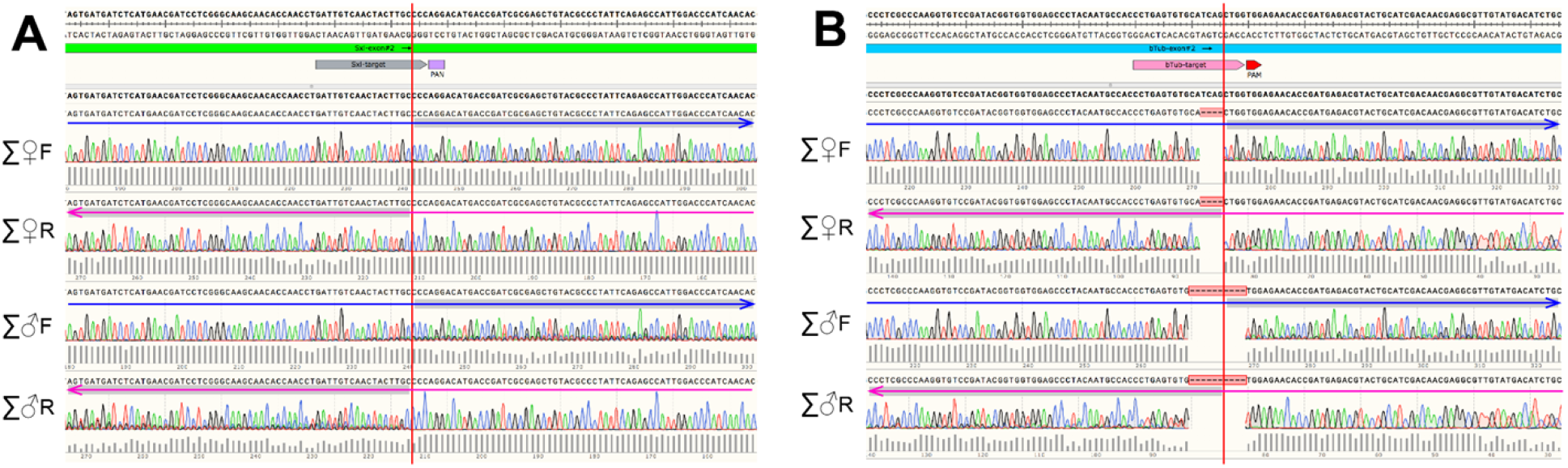
Basal *Hsp70Bb-Cas9* expression in somatic tissues of *TI-pg^sxl,βTub,Hsp-Cas9^* flies. The *Drosophila heat-shock protein 70B* (*Hsp70Bb*) promoter is known to drive a baseline expression at 25°C^28–30^. To assess whether *Hsp70Bb-Cas9* is expressed at 18°C, we sequenced target sites in *sxl* (**A**) and *βTub* (**B**) using the DNA extracted from multiple females (∑♀) and males (∑♂) of *TI-pgSIT^sxl,βTub,Hsp-Cas9^*/+ flies reared at 20°C. The presence of induced *indel* alleles at the cut site (indicated by red lines) among *wt* alleles for both *sxl* and *βTub* loci causes ambiguity (multiple peaks) of sequence reads. Directions of sequence reads are indicated with blue (forward primer) and red (reverse primer) errors.

